# Distorting anatomy to test MEG models and metrics

**DOI:** 10.1101/2025.04.03.646622

**Authors:** José David López, Yael Balbastre, John Ashburner, James J. Bonaiuto, Gareth Barnes

## Abstract

Current flow that gives rise to non-invasive Magnetoencephalographic (MEG) data derives predominantly from pyramidal neurons oriented orthogonal to the cortical surface. The estimate of current flow based on extra-cranial magnetic fields is a well-known ill-posed problem; however, this current distribution must depend on anatomy. In other words, a veridical estimate of current flow should discriminate between true and distorted versions of the brain. Here, we make use of advances in diffeomorphic brain shape modelling to construct a set of parametrically deformable cortical surfaces. We use a latent space of 100 components to construct cortical surfaces that are representative of the population. We show how these geometric distortions can be used to quantify the performance of MEG source reconstruction algorithms and metrics of fit.

## Introduction

Magnetoencephalographic (MEG) and electroencephalographic (EEG) recordings enable reconstructions of spatio-temporal images of human brain function. Estimating these images from data measured outside the head requires additional assumptions, which are continually being refined, tested, and updated. We have proposed that anatomy (available from non-invasive Magnetic Resonance Imaging – MRI) provides a valuable ground truth against which to decide between assumption sets (Barnes et al., 2006; Little et al., 2018; López et al., 2017; Stevenson et al., 2014). We know that the primary current flow (which gives rise to the external magnetic fields) is bound (in the cell bodies of pyramidal neurons) to anatomy. Terefore, imprecise or distorted anatomical models must lead to sub-optimal functional estimates. We hypothesize that more accurate current flow models will be more likely given the correct, rather than incorrect, anatomy.

The main issue addressed by this paper is the generation of realistic (brain-like) surrogate anatomy. (Barnes et al., 2006) used rotated versions of the true anatomy within a spherical volume. (Stevenson et al., 2014) and (Little et al., 2018) progressively removed higher-order components from spherical harmonic approximations to the cortical manifold. While (López et al., 2017) produced surrogate brains from linear mixtures of these harmonic components. All these cases, whilst providing proof of principle, set a relatively low bar on algorithm performance as the surrogate brains were not anatomically realistic.

In contrast, (Troebinger et al., 2014) found higher evidence for generative models (of the MEG data) based on an individual’s true cortical anatomy, rather than another person’s brain warped into the same individual’s head. Here we pursue this direction of using realistic surrogates but take advantage of recent advances in diffeomorphic brain shape modelling (Ashburner et al., 2019). This work was originally developed to enhance data mining of large MRI datasets, by parameterizing each structural MRI into a relatively small number of latent variables. Importantly, these latent variables, which describe a population based on large structural databases, are Gaussian. This means that it is possible to describe an individual’s brain shape as a vector of values and construct perturbations of this vector that distort this brain within realistic limits. The deformations here are subtler than one might expect when exchanging brains (in which patterns of gyrification may change), as the individual topology is preserved.

The M/EEG source reconstruction problem is an ill-posed model inversion, so it must be constrained by imposing biologically and physically plausible assumptions. Different M/EEG source reconstruction methods entail different assumptions about the underlying current distribution, yet all produce subjectively plausible current estimates. This work aims to develop an objective pathway that can be used to compare any part of the M/EEG analysis pathway. Here we compare among different source reconstruction schemes (and their underlying assumptions) and goodness of fit metrics.

## Methods

### Diffeomorphic modelling

The diffeomorphic algorithms used in this paper have been described extensively elsewhere (Balbastre et al., 2018). We briefly summarize them below.

Diffeomorphic shape modelling assumes that any individual anatomy can be described by a diffeomorphic (i.e., smooth and invertible) transform applied to a common template (Miller et al., 1993). Conversely, given a collection of observed anatomies, an empirical template encoding the average anatomy of a population can be obtained by minimising its distance to all elements of the collection in a procedure known as geodesic shooting (Ashburner & Friston, 2011; Miller et al., 2006). In this framework, diffeomorphisms are encoded by an “initial velocity” and the distance between anatomies is penalised on the squared gradients (or other differential features that describe smoothness) of this velocity field. After template construction, the posterior distribution of velocity fields can be efficiently estimated by performing a Bayesian residual component analysis (Bishop, 1999; Kalaitzis & Lawrence, 2011), a variant of principal component analysis that allows ‘residual’ variance (here, known smoothness) to be factored out of the projected variance when determining principal modes of variation. This analysis allows each velocity field (and, therefore, each anatomy) to be described by a small number of latent projections (Ashburner et al., 2019; Balbastre et al., 2018; Zhang & Thomas Fletcher, 2015). New anatomies can then be encoded in this low-dimensional latent space by registering them with the template and projecting the resulting velocities onto the principal basis. Furthermore, random anatomies can be sampled from the learnt model by sampling latent codes from the standard Normal distribution.

Here, 662 structural MRI scans (with a 1:1 male/female ratio) were randomly sampled from four large datasets (“IXI Dataset”; “OASIS”; “COBRE”; “ABIDE”). They were segmented (Ashburner & Friston, 2005), resliced to 1.5 mm, and the grey and white matter tissues were diffeomorphically aligned with their common mean (Ashburner & Friston, 2011). A 100-component Bayesian residual model was fitted to the resulting velocity fields. Brains were rigidly aligned, meaning that brain size was encoded within at least one of these modes.

### Parameter space

The shape model encodes variability using 100 eigenvectors to account for normal brain shape variation. Each individual therefore has a vector (**z**) of 100 values that describe their brain shape and a corresponding cortical surface mesh. We then distorted this surface mesh by manipulating this individual vector (**z**). In order to avoid any confounds due to the volume of the mesh changing (i.e. sources becoming nearer to the sensors), we first modulated different ranges of components (e.g. 1-100, 4-100 etc) and selected a range (8-100) that maintained an approximately constant brain volume (figure S1). We sampled each of these 93 components across a range of fixed points, which traversed a range of z-scores with that closest to true brain at the centre (figure 1A). We define a distortion trajectory as moving from either {−*Z*, +*Z*} or {+*Z*, −*Z*} (randomly assigned per component *i*) as index *j* moves from 1 to *N*. The value of each component (*i*) on the distortion trajectory is given by *d*_*i,j*_. The random selection of direction per component (determined by the random seed that initializes the trajectory) means that it is possible to create several close to orthogonal trajectories for each subject (Figure S2). In other words, for each subject it is possible to create surrogate cortices that are distant to one another yet equally displaced (determined by *j*) from the true cortex (Figure 1B).

**Figure 1.**
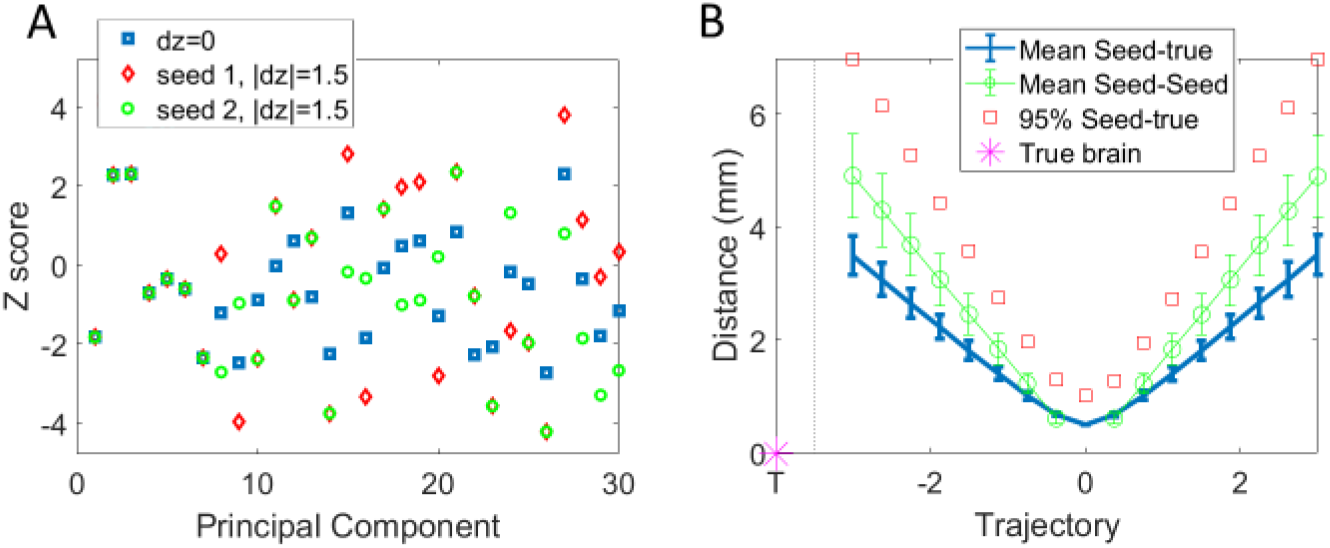
**A)** Example of the first 30 (of 100) components defining the true cortical surface **z** (δz = 0, blue squares) for an individual. In addition, two trajectories (based on different random seeds) are shown for |δz| = 1.5. Note that only components 8 and beyond are distorted. **B)** Mean and standard deviation vertex-vertex distances between cortices on eight trajectories (eight random number seeds) and the true surface (blue solid). Also shown are the mean 95^th^ percentiles (red-squares) of vertex-vertex distances (i.e. 95% of the surface vertices are closer than this). An additional point (asterisk with x-label ‘T’’), at mean distance zero, has been added to represent the true (undistorted) cortical surface for consistency with later figures. Mean and standard deviation trajectory-trajectory (or seed-seed) vertex-vertex distances are shown as green-circles.

Formally, we define a random seed which gives a deterministic vector of random signs *s*, where

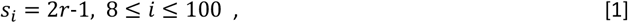

where *r* is drawn from a random binary distribution *P*(*r* = *1*) = 0.5, *P*(*r* = 0) = 0.5, and *s*_*i*_ is therefore (pseudo-randomly) either +1 or -1.

For higher order components this is

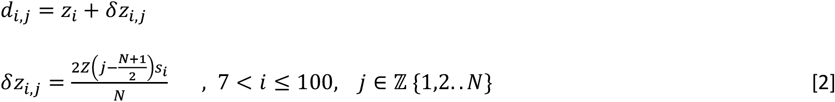

Whereas for the lower order components (determining brain size etc.)

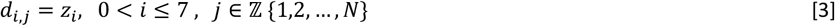

where *z*_*i*_ is the true eigenvalue representing the individual’s brain at component *i*. The amount of distortion is given by integer *j*, which moves *d*_*i,j*_ across a range from +*Z* to −*Z* or (vice versa depending on *s*_*i*_). *N* is an odd integer defining the number of cortices within a trajectory. In this case, *N* = 17, *Z* = 3.

Note that the point 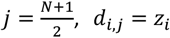, on the trajectory corresponds to the best approximation of the true surface. Each individual trajectory is defined by a random seed that determines the pseudo-random sequence (of +1 and -1) in **s**.

The individual cortical surface was constructed using FreeSurfer (Fischl, 2012). We used a white matter cortical surface model consisting of ∽30,000 vertices (current dipoles) with orientation defined by the local surface normal. Each new surface gives rise to a new source model defining a set of current dipoles at each mesh vertex location and orientation. The extra-cranial field due to this primary current flow in turn depends on the intervening media. Here we made use of a single-shell model (Nolte, 2003) to provide a 10^th^ order multiple spherical harmonic approximation to the inner-skull boundary. That is, the exploration of parameter space is an exploration of different source models (each defined by a different surface mesh).

We also constructed a null forward model in which the correspondence between the MEG sensor data and the forward model predictions were removed. We did this by permuting the channel data with respect to the sensitivity (or lead-field) matrix linking the field produced by any single dipole to the sensor array; i.e. the data remained the same, but the forward model was now meaningless.

### Model comparison

The next step is to assess how well our MEG methods discriminate between the true and surrogate anatomies. A practical metric of model fit will be one that is maximized when the underlying anatomy (by which the current flow is bounded) is correct (Barnes et al., 2006; Little et al., 2018; López et al., 2012, 2017; Stevenson et al., 2014).

In this work, we assess three goodness of fit metrics:

- **Negative variational Free energy** (*F*) (Friston et al., 2007) (also known as the evidence lower bound - ELBO).
- **Variance explained** (i.e. the squared correlation coefficient between measured and predicted data).
- **Cross-validation error**. (Bonaiuto, Rossiter, et al., 2018; Browne, 2000; Little et al., 2018) Here we compute cross-validation error as the average root mean square difference in femto-Tesla (over the 10 cross-validation runs) between the predicted and measured values in the 10% of MEG channels excluded from the source reconstruction (and randomly selected in each cross-validation run).

We use common inversion algorithms to explain the measured MEG data as current flow constrained to each surface on the trajectory in turn. Each algorithm can be defined by the modelling assumptions that determine the covariance of neuronal current flow across the cortex (López et al., 2014; Mosher et al., 2003). The models tested were all implemented in the open-source software package SPM (Litvak et al., 2011), and were:

- **IID**: Minimum Norm algorithm (Hamalainen et al., 1993).
- **GS**: Greedy Search Algorithm - enforcing sparseness (Friston et al., 2008).
- **EBB**: Empirical Bayes Beamformer (Belardinelli et al., 2012),

### MEG empirical data

We used previously published data from MEG recordings of eight subjects using head-casts (Bonaiuto, Meyer, et al., 2018; Little et al., 2018). See data-availability section for an updated link. The head-cast minimizes subject movement during the experiment and gives a precise estimate of the relative positions of the sensor array and head (Meyer et al., 2017; Troebinger, López et al., 2014). In brief, recordings were made using a 275-channel CTF Omega MEG system. The data were sampled at 1200 Hz, filtered (5th order Butterworth band-pass filter: 2–100 Hz, Notch filter: 50 Hz), and down-sampled to 250 Hz. Eye-blink artefacts were removed using multiple source eye correction (Berg & Scherg, 1994). Participants completed a visually cued action decision-making task in which they responded to a Random Dot Kinematogram (RDK) (displayed for 2 s with coherent motion, either to the left or right). Following a 500ms delay, an instruction cue appeared, pointing either to the left or the right, and participants were instructed to press the corresponding button (left or right) as quickly and as accurately as possible.

The processing of the data was either focused on trials locked to the onset of the random dots, the instruction cue, or to the button press.

## Results

We now move through the space of possible cortical surfaces and examine the impact on MEG reconstruction algorithms and fit metrics. We first show how the cortex is progressively deformed from the anatomical ground truth. We go on to test whether this deformation has any influence on the metric or method of source reconstruction. The logic is that distortions from the true anatomy will be of little consequence for sub-optimal metrics or reconstruction assumptions. We initially focus on a single dataset and then expand to the group of datasets (and distortion trajectories).

### Deformations

Figure 1A shows two potential distortion trajectories (Eqs. 1 and 2). The shape of any brain in the population can be defined by a 100-element vector. This subject’s brain is defined by vector **z** (blue squares). We then define a random deformation trajectory to move each person’s brain about **z** in 17 successive steps. This produced 17 cortical surfaces, one for each trajectory step (see Figure 1B). To illustrate the symmetry, we plot the trajectory as moving from −3 to +3; although at each component this direction will be randomly selected (Eq. 1). Surfaces 1 and 17 are at the extremes of the trajectory, with surface 9 at the origin, where δ*z* = 0 (i.e. closest to the original cortical surface). In addition, the true cortical surface (labelled ‘*T*’) on the x-axis is added for reference (as the trajectories do not capture all of the anatomical variance). Figure 1B shows the average distance of each vertex to the corresponding vertex on the original cortical surface *T* (distributions shown in Figure S1). Also shown in Figure 1B are the distances between surfaces lying on different trajectories (determined by random seeds which set the pseudo-random signs in ***s***) and one another. That is, even though the trajectories are all a similar distance from the true brain; they are also distant from one another. Figure S2 shows the correlation (mean r^2^=0.046) between the vectors ***δz*** that define each trajectory.

Figure 2 shows three points on one trajectory of brain deformations, alongside similarly deformed scalp surfaces. The dotted circle highlights one of the regions affected by the deformation. As it is difficult to spot these cortical deformations by eye, the subject’s scalp surface is shown below to illustrate the deformation as |δ*z*| increases in magnitude. Note the gradual distortion of the face moving from left (truth) to right (distortion of lips and face).

**Figure 2.**
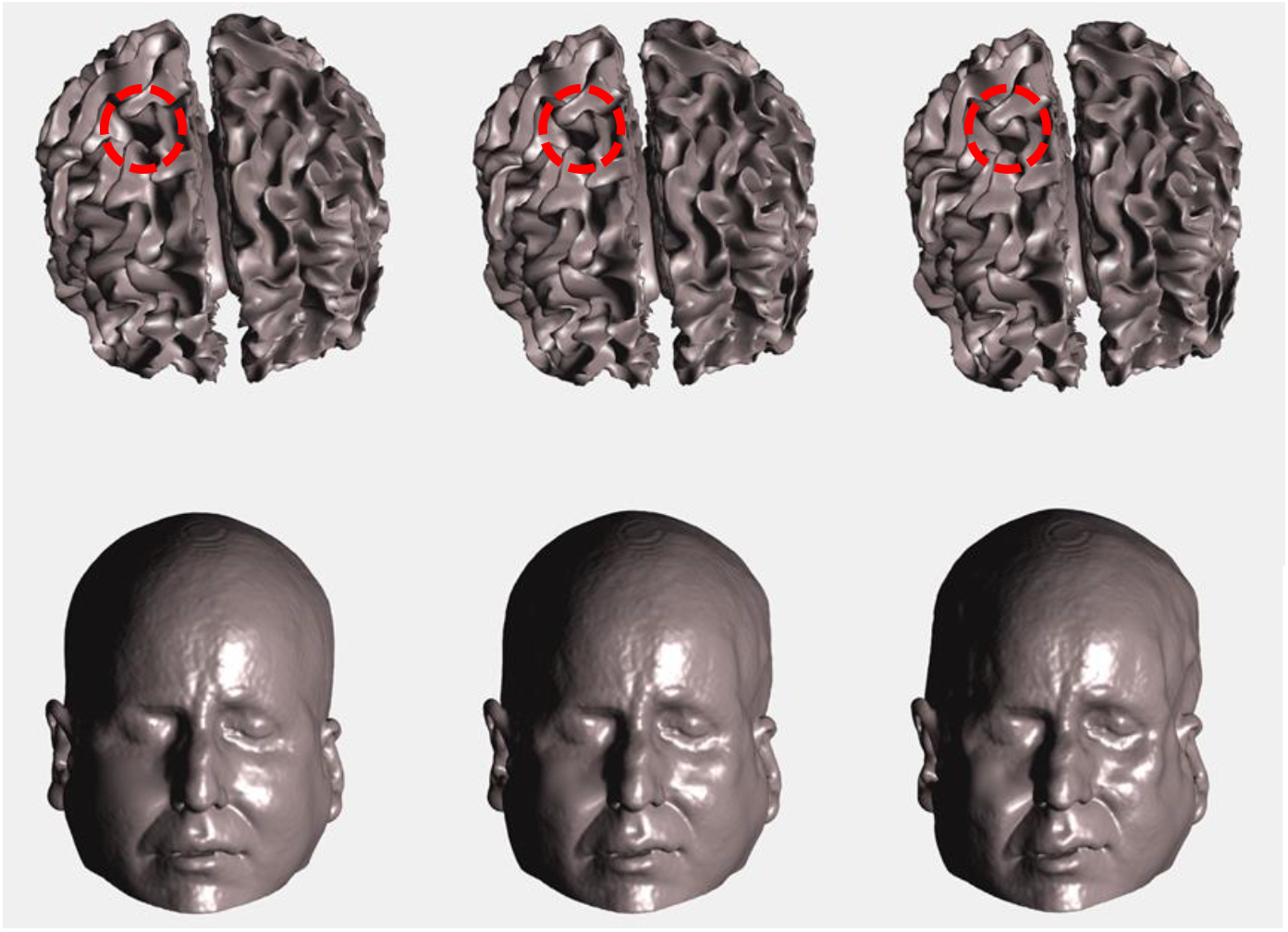
Top panel. Cortical meshes from one example trajectory at points |δz| = 0, |δz| = 1.5 and |δz| = 3 (left-right respectively), which show increasing amounts of distortion. The red-dotted circle highlights one of the narrowing of one of the sulci during the distortion. For illustration only, the lower panel shows the (70% reduced in size) scalp surface of this subject distorted in the same way.

### Goodness of fit metrics

We now examine (Figure 3) different metrics of model fit and different inversion models as the anatomy moves through this deformation trajectory. In an ideal scenario, one would hope that the model fit metric (based purely on the MEG data) improves for brain structures closer to the truth. If the model fit does not perform in this way, it could be a poor metric, or the underlying algorithm, data, or forward model might be sub-optimal.

**Figure 3.**
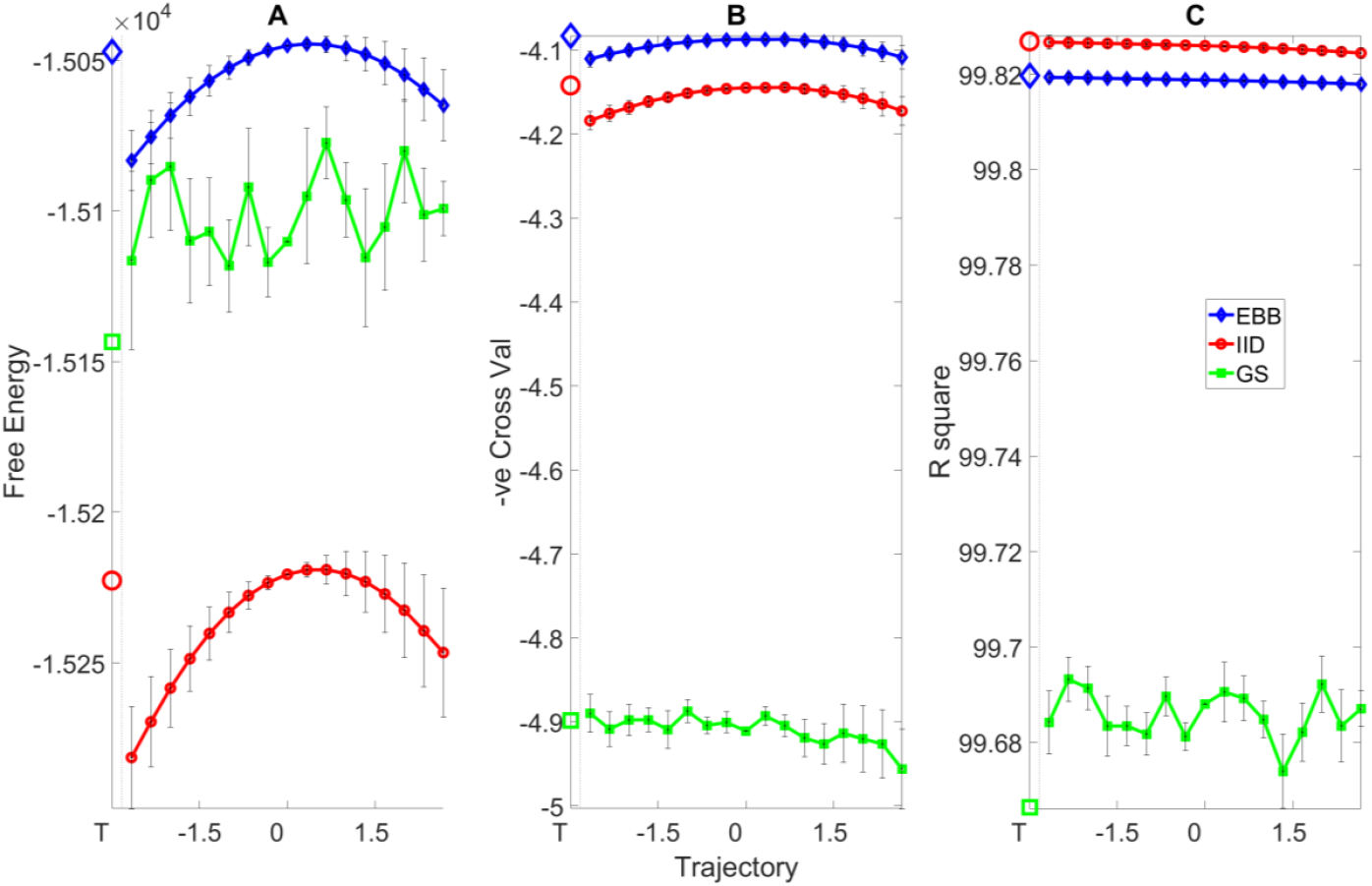
EBB (blue diamonds), IID (red circles) and GS (green squares) inversion algorithms with Free energy (A), cross-validation (B) and R square (C) cost-functions for different distortions of the cortical mesh. The cortical mesh moves from the true mesh (T) through a distortion trajectory at each component (i) beginning and ending at |δz| = 3 passing through δz = 0. The two ends of the range are the same distance from |δz| = 0 (Figure 2B) but in different directions (δz_i,1_ = −δz_i,N_). **A)** Mean and standard error (black bars) of Free energy metrics for the three inversion algorithms across eight trajectories for a single subject. **B)** Mean and standard error of (negated) cross-validation errors for the same trajectories and data as in A. **C)** The R square statistic, or percent variance explained, for the same trajectories and subject data as in A.

In Figure 3, all metrics are plotted such that the most likely model has the highest value (i.e., cross-validation error is negated). Panel A shows the negative variational Free Energy for three different models of underlying current flow expressed on a cortex that is systematically distorted. Free energy scores for EBB and IID peak close to the true anatomy (i.e. at point T and around δ*z* = 0); GS, however, is rated as more likely model (higher Free Energy) than IID yet it does not show any sensitivity to the varying anatomy. Panel B shows fits to the same data as rated through 10-fold cross-validation error. Once again EBB and IID have maximal scores around the true brain (T) and smaller distortions; in this case, GS also does not co-vary with anatomical variation but it is also rated the poorest model. Panel C shows the R square metric of model fit. It is clear that this fit metric does not discriminate between veridical and erroneous anatomical information. This is most likely because there is no penalty for over-complex solutions (much like a regression equation in which R square increases monotonically with parameters).

The previous results relate to evoked response data for subject 1, condition 1 across eight distortion trajectories (eight random seeds). Figure 4 shows the summary ranked Free Energy values across eight subjects with three conditions and eight trajectories per condition per subject (i.e. 8 × 3 × 8 = 192 trajectories). The ranking removes the relative scaling across runs and participants and allows one to summarize the model score from most (18) to least (1) likely.

**Figure 4.**
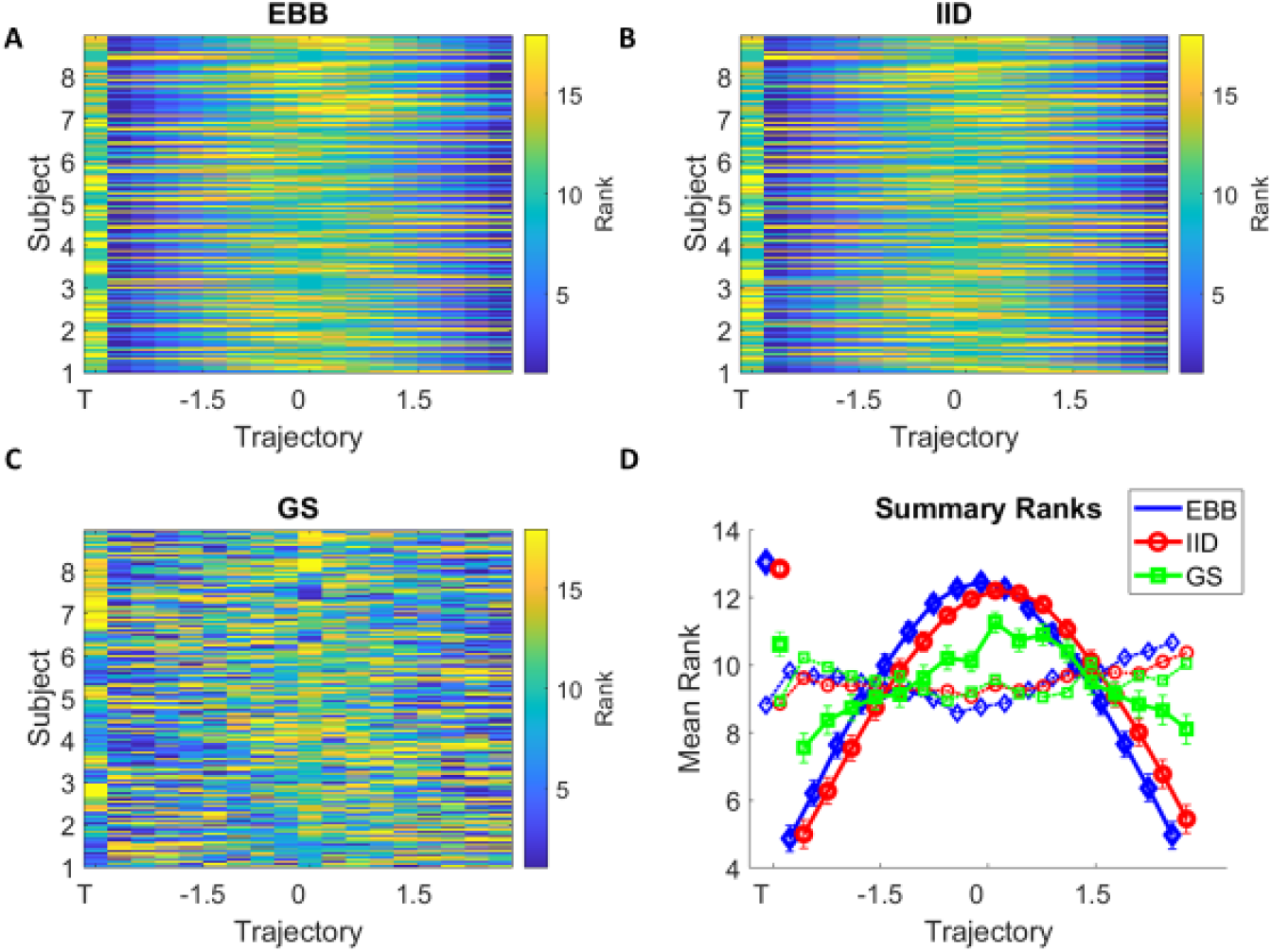
Group results for eight subjects, three conditions, and eight trajectories. Panels A-C correspond to the EBB, IID and GS algorithms respectively. Each row is a rank-transformed (highest value has maximum rank) set of 18 (one true surface plus 17 surfaces on the trajectory) Free-Energy values. High-ranks near the true brain (T) or close to zero distortion indicate that the algorithm is selective of the true anatomy. Panel D is the summary plot of panels A-C. Average (and standard error) ranks for EBB (blue diamonds), IID (red circles) and GS (green squares) are shown. The dotted lines show the corresponding distributions using the null forward model (corresponding panels A-C in Figure S4). Curves are staggered along the x-axis for display purposes.

Figures 4A-C show the rank-transformed curves per subject, condition and trajectory for EBB, IID and GS algorithms respectively. Each row is a trajectory and larger ranks imply more likely models. Higher ranks clustered towards the centre of the trajectory or around the true anatomy (T) suggest that the model fit is sensitive to true anatomy. Figure 4D summarizes panels A-C in terms of mean and standard error of the rankings for the different inversion schemes.

These curves can be compared based on how much they differ to ideal rankings (i.e. the true cortex has maximal rank, the most distorted cortices have the lowest ranks). For each row of the matrices in Panels A-C, one can compute the absolute rank differences for each of the algorithms from ideal (Figure 5A). Algorithms more sensitive to the true anatomy will have lower rank differences to ideal performance. This can be compared to the performance using the permuted lead-fields (black-dotted). In this case, all three algorithms make a significantly improved estimate of anatomy using the true rather than null (permuted lead-fields) model (figure 5A).

**Figure 5.**
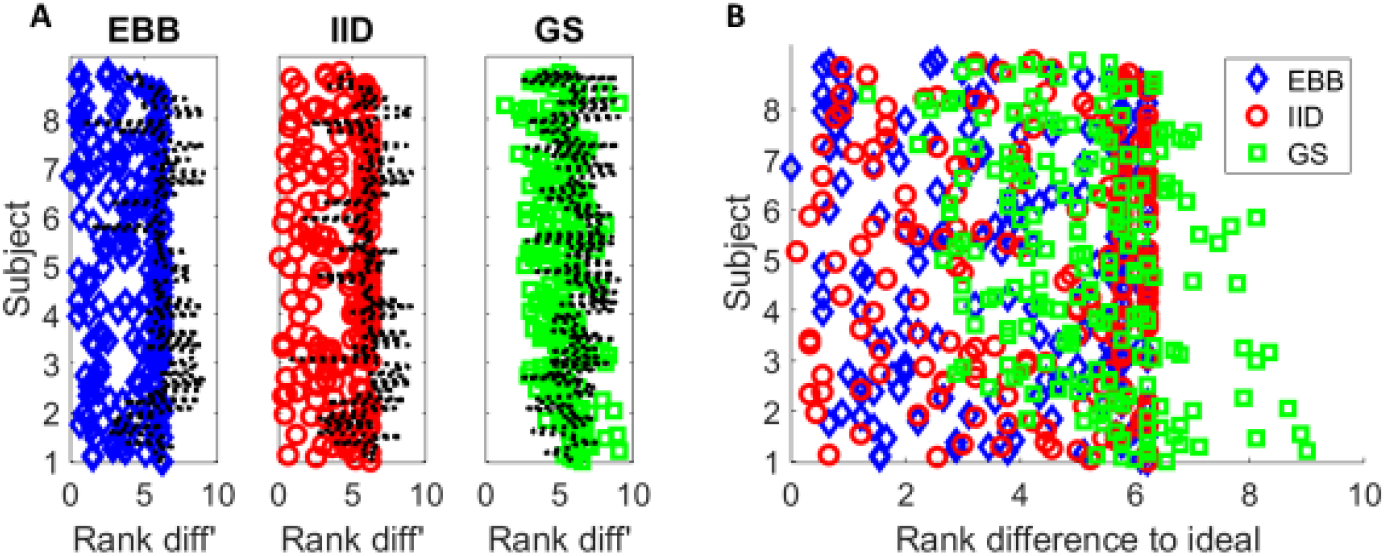
**A)** Mean absolute rank difference to ideal for EBB (blue diamonds), IID (red circles) and GS (green squares) algorithms. Within each panel, each row corresponds to a dataset/trajectory in Figure 4A-C. The black dotted lines show the performance of the algorithm with permuted lead-fields. Smaller rank differences mean that the fit metric (Free Energy) increases for surfaces closer to the true brain. All three algorithms significantly improve on the estimate of true anatomy (p<2.6×10^−27^, t=-12.7, df=191; p<7.4×10^−21^; t=-10.57, df=191; p<4.7 x10^−7^, t=-5.22, df=191 for EBB, IID and GS respectively) when using real (symbols as in 5A) as compared to permuted data (black dotted). **B)** Mean absolute rank differences (to ideal performance) between algorithms. Both EBB and IID algorithms significantly improve over GS (p<1.72×10^−12^, t=1.42, df=191; p<9.95×10^−10^, t=6.43, df=191) but there was no significant difference between EBB and IID performance (p=0.159, t=1.42, df=191).

Comparing between algorithms, EBB and IID depend (significantly) more on the true anatomy than GS (figure 5B) but no significant difference between EBB and IID was found.

## Discussion

We leverage the knowledge of true cortical anatomy to compare among MEG analysis pathways. We have done this using realistic and quantifiable deformations to distort an individual’s cortical surface. We show that realistic deformations of anatomy can be used to compare among different models and metrics of model fit. Importantly, performance can be quantified objectively in terms of a distance from ground-truth anatomy. In this study, we found that fit metrics, such as Free Energy and cross-validation accuracy, are optimized when the anatomy is correct. We also found that inversion schemes based on the assumption sets behind IID and EBB were most sensitive to the true anatomy.

Here we examine inversion methods and fit metrics but the approach could be used to empirically determine best practice at any stage of the MEG (or EEG) pathway from the choice of forward models to sensor design. The three algorithms used are available in the current SPM release.

It should be noted that the implementation of GS used here is most likely sub-optimal in that only a fixed number of discrete (pre-defined) cortical patches are allowed to be active. Previous work has combined the greedy-search with an automatic relevance determination (ARD) stage (Friston et al., 2008): the greedy search to find variance components (source distributions) that maximize the model evidence and ARD to prune away those that make least contribution to the evidence. Alternatively, more computationally intensive approaches have been suggested in which model evidence is assessed over multiple possible patch locations to overcome problems with sparse sampling; and this in turn is run over multiple iterations to avoid local maxima (Lopez et al., 2012). In this study, GS is a useful vehicle to show that sometimes there may be a disconnect between the mathematically most likely model of the data and the true generative model bound to the anatomy (Figure 3A).

We had not expected to find so little difference between the sensitivity to anatomy of EBB and IID inversions. There could be a number of reasons for this: i) The activity was primarily cortical and superficial and so it could be more difficult to distinguish between IID (typically superficially biased) and EBB estimates. ii) The EBB is not a classic beamformer algorithm (Van Veen et al., 1997) but rather uses an empirical beamformer prior as part of a generative model (Belardinelli et al., 2012). In cases in which a conventional beamformer might fail (for example zero lag correlated and distinct sources) there is no useful information in the prior and it becomes flat – defaulting to minimum norm behaviour. In other words, EBB and IID will tend to the same algorithm in certain cases. iii) Finally, and perhaps most-likely, neither the assumptions between EBB or IID generative models match the true neuronal generative model and/or algorithm performance is constrained by some other factor (for example sub-optimal volume conductor model).

There are several key limitations here. The cortical deformations are based on eight (random) distortion trajectories for all subjects. In order to observe any change in how the measured MEG data can be expressed on the cortical surface, we rely on the distortion perturbing cortical sources that will influence the estimate of current flow. However, distortion trajectories must exist that minimally influence this estimate; in which case we would see no change in model fit with distortion. Using longer-recordings and novel data processing techniques (Woolrich et al., 2013) would allow one to extract multiple useful stationary modes from these data (Little et al., 2018; Martinez-Vargas et al., 2016)

The averaged evoked data shown are specific to one experiment derived from three time-windows and in a single frequency band for a single dataset. We cannot therefore generalize findings across other datasets or to data due to induced (rather than average-evoked) changes. We used the white matter (rather than pial) surface and made no attempt to optimise the orientation of current flow (Bonaiuto et al., 2020). We used exclusively cortical surfaces; but if the manifold on which sub-cortical sources sit is well defined (Attal & Schwartz, 2013; Meyer, Rossiter, et al., 2017) the same approach could be used. Likewise, rather than distort the cortical surface, the same code can be used to also distort any of the conductivity boundaries (such as the inner skull). Finally, the error from the true brain to its representation in 100 principal components was relatively large (∽0.5 mm). Further work could explore parameterizable representations of the cortical surface which come closer to zero error and allow for finer grained distinction between methods.

Note that these data are based on head-cast recordings. Here, subjects wore individualized foam helmets that fit the scalp internally and the MEG system externally. Therefore, the relative location of the subject’s anatomy and the sensors is well-defined and fixed. That is, the subjects’ heads were immobilised in the scanner. This scanning method is time-intensive, only appropriate for healthy compliant subjects, and carries a certain amount of risk (due to the potential of MEG system movement whilst the subject’s head is fixed). There is now, however, a new generation of MEG sensors known as optically pumped magnetometers (OPMs) (Tierney et al., 2019), which can be worn for long periods, and their location relative to the underlying anatomy can be well defined. In other words, although head-cast recording may be limited, OPMs provide an exciting and expanding field to test these methods further. In the limit of increasing measured data for an individual, these tests between competing anatomies should converge to a single solution (the true individual anatomy). The more data available per-subject the more constrained (and well-posed) the problem becomes.

## Supporting information

Supplemental for Lopez et al.

## Ethics

We used previously published data from MEG recordings of eight subjects using head-casts (Bonaiuto, Meyer, et al., 2018; Little et al., 2018). The study protocol was in full accordance with the Declaration of Helsinki, and all participants gave written informed consent after being fully informed about the purpose of the study. The study protocol, participant information, and form of consent, were approved by the UCL Research Ethics Committee (reference number 5833/001).

## Author Contributions

GRB, JDL, YB-conception. GRB, JDL, JB-MEG specific code. YB, JA-Structure specific code.

## Data and code availability

The code is part of the current development version of SPM (https://github.com/spm/spm).

The deformation templates will be automatically downloaded the first time the algorithm is used.

The MEG evoked response data, individual MRIs and cortical surfaces are available here https://figshare.com/s/e3ff829753a76bcfda22

The additional code to reproduce Figure 3 (and hence all other figures) can be found at https://github.com/barnesgr123/spm_distort/

The complete dataset, originally from (Bonaiuto, Meyer, et al., 2018; Little et al., 2018), can be found at https://osf.io/eu6nx.

## Acknowledgements

JDL Was funded by the Colombian MinCiencias project contract 1222-852-69927.

YB was funded in part by the MRC and Spinal Research Charity through the ERA-NET Neuron joint call (MR/R000050/1), and in part by a fellowship from the Royal Society (NIF\R1\232460).

This research was supported by the Discovery Research Platform for Naturalistic Neuroimaging funded by the Wellcome (226793/Z/22/Z). We would like to thank the UCL MEG group for their helpful and constructive comments.

