## Supplemental for Lopez et al. for "Distorting anatomy to test MEG models and metrics"

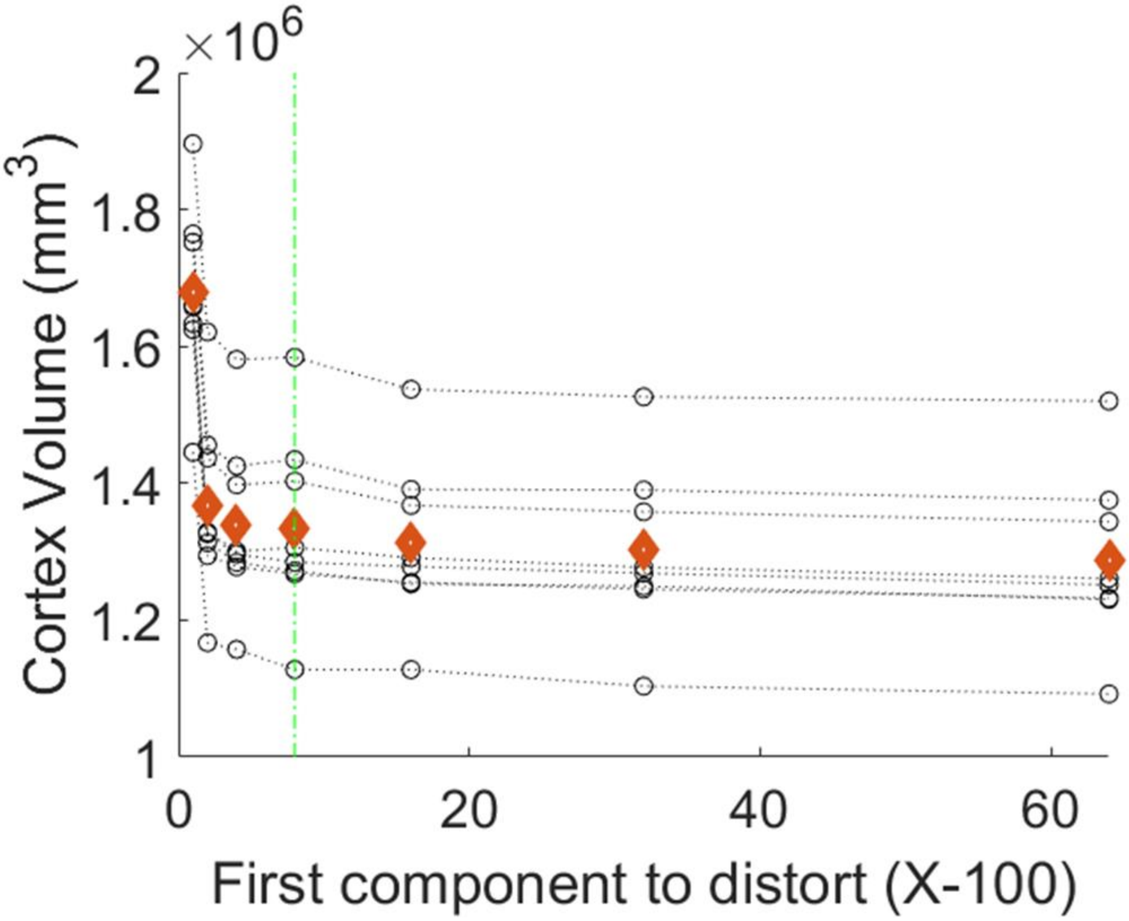


**Figure S1.**

Cortical volume (circles) for 8 subjects as a function of the principal components distorted (1-100, 2-100 etc). Red diamonds show the mean volume changes. Green dotted line highlights range (8-100).

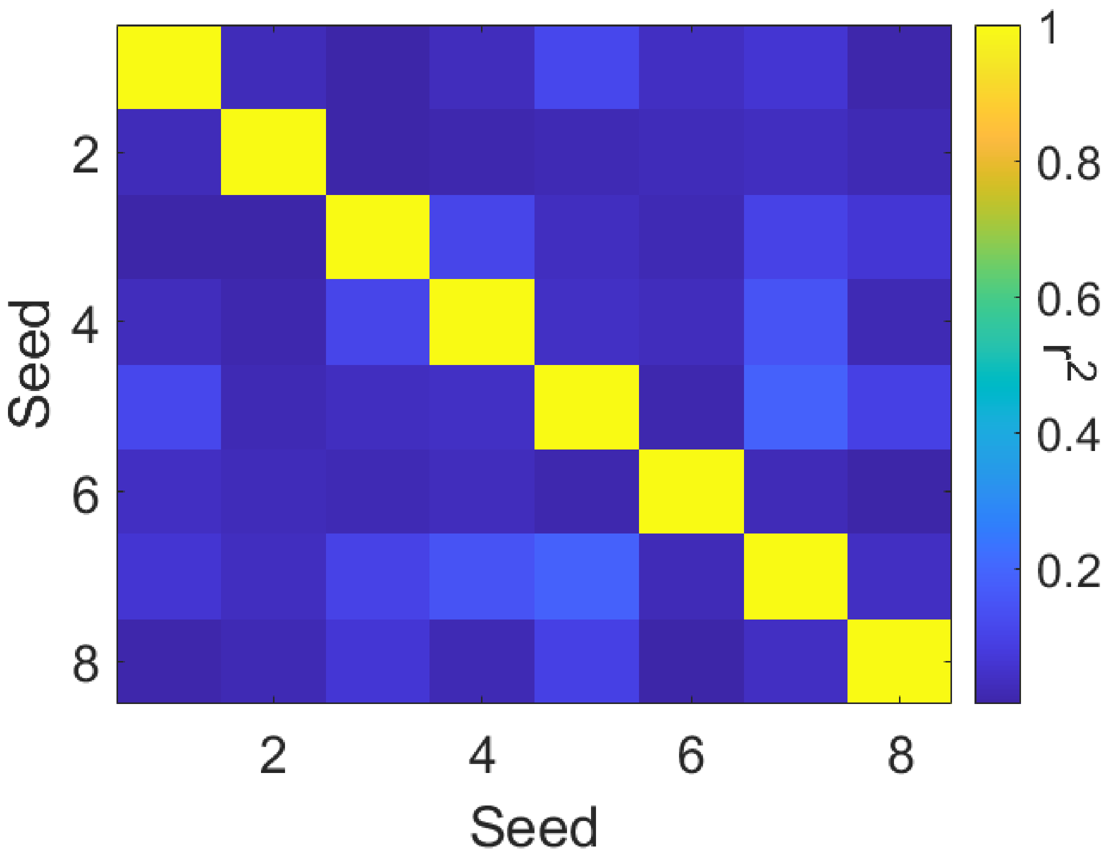


**Figure S2**

The correlation between the 100 element PCA z-score vectors defining the 8 different distortion trajectories (each defined by a random seed) used.

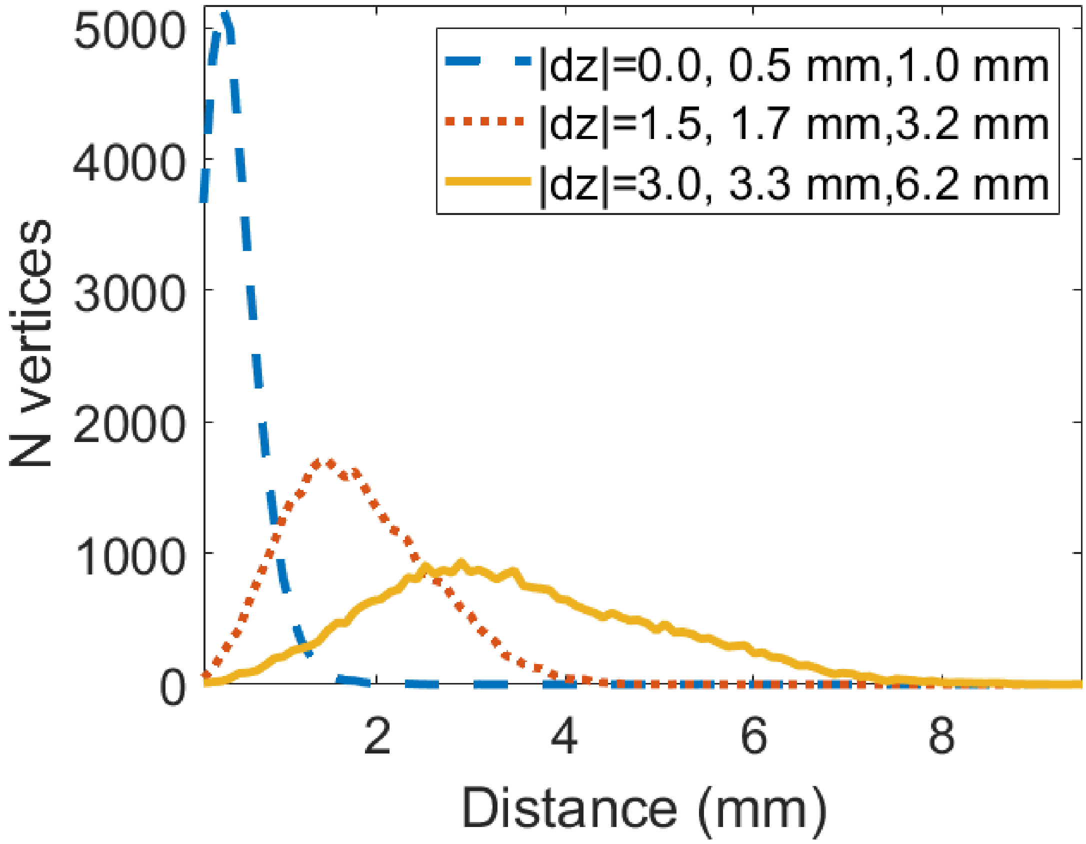


**Figure S3**

The distribution of vertex-vertex distances (between the true and distorted cortex) for different points |dz| along the distortion trajectory. The legend shows the value of |dz|, the mean vertex-vertex distance and the 95$^{th}$ percentile of vertex-vertex distances respectively.

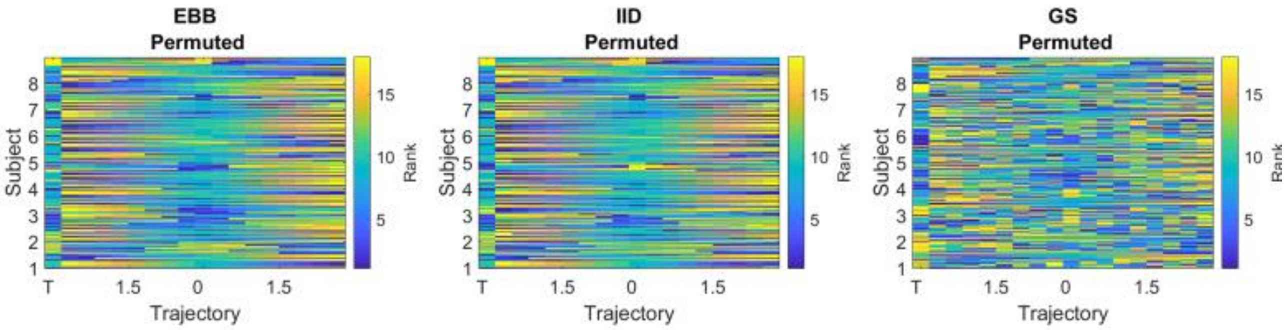


**Figure S4**

The rank scores analogous to figure 4 in manuscript but using the null (permuted lead-field) forward model. The means and standard errors of these distributions are shown in panel 4D of manuscript as dotted lines.
